# Sleep cycle of a fin whale based on reprocessing of old satellite observations

**DOI:** 10.1101/655886

**Authors:** Denise Viale, Laurent Koechlin, Carli Viale

## Abstract

Reprocessing of old data from the satellite tracking of a fin whale in 1991 allowed us to detect new loops on its path and to show the significance of these loops. Originally, five loops on five consecutive days had been detected. The reprocessed data reveals at least 21 loops, plus many other phases of inertia, together with additional information on the whale’s position at the surface during these phases of inertia. Based on these new interpretations, we deduce a sleep cycle for this animal.

**Résumé:** Mise en évidence à partir d’un suivi satellitaire d’une phase de sommeil quotidienne chez un baleinoptère.

La fixation d’une balise Argos sur un rorqual (*Balaenoptera physalus*) donna lieu à un suivi satellitaire de septembre à novembre 1991, qui visait à démontrer notre hypothèse sur leurs déplacements migratoires. Notre suivi n’avait pas été assez long pour déterminer si le baleinoptère marqué sortait de Mediterranee mais il avait permis de mettre en évidence quelques phases d’inertie dans la vie de la baleine, identiques à celles observées dans les trajectoires de bouées dérivantes utilisées en océanographie physique.

Nous avons repris ces anciennes données avec de nouveaux traitements. Outre la mise en évidence de l’adéquation de la trajectoire et du comportement de l’animal avec les caractéristiques hydrologiques et trophiques de chaque localisation, cela a permis de détecter une vingtaine de nouvelles boucles et des phases d’inertie dans la trajectoire de la baleine, et de les apparenter à des phases de sommeil. Pour cela il fallait démontrer quélles se répétaient chaque jour.

## 1. Introduction

In the late 1980s, the Argos technology provided for the first time efficient means of tracking cetaceans by satellite, at scales suited in space and time to their migrations.

The initial conception of our “Argocet” tracking program dates from 1986. Its development phase, from 1988 to 1991, led to a dedicated Argos platform transmitter (Argos PTT). This small device provided tracking data from 22 September to 2 November, 1991, attached to a fin whale (*Balaenoptera physalus* Linnaeus, 1758 (*Balaenopteridae, Mysticeta*), abbreviated “Bp” hereafter).

Our tracking program was intended to demonstrate our hypothesis on the migratory course of fin whales. Bp are present during winter and summer in the Mediterranean sea [1] [2] [3], but these two populations are probably different [4]: one, present in winter, migrates in summer towards the North Atlantic [5], whereas the other, more abundant in summer, migrates to the South Atlantic according to our hypothesis. The tracking record did not last long enough to determine whether ‘our’ whale eventually left the Mediterranean, but the loops in its recorded track revealed inertial phases in its life cycle [6] [7] [8].

These loops of variable size (a few km) are similar as those observed for drifting buoys used in oceanography [9] [10]. However, linking these loops to sleep phases requires observation of a repeated occurrence over many days, and only five loops during five consecutive days had been detected in the initial study [11].

We reprocessed these old data with modern means. This led to the detection of at least 21 loops in the whale’s track, in addition to other phases of inertia, and allowed correlations to be made with the hydrological, trophic and meteorological conditions at the different points.

## 2. Equipment and Methods

When the tracking data was collected in 1991, D. Viale and collaborators only had the old Argos technology available. Now, the new Argos system and the global network satellite system (GNSS) are much more powerful. However, few large cetaceans have been tracked since, and the original data can still be exploited.

The Argos buoy used for this survey, conceived in Nice by J.J. Pesando, withstands speeds up to 23 knots and depths to 600 m. This 2.5 kg transmitting platform (PTT) consists of a carbon fibre container housing the emitter, covered with polyurethane. It contains lithium batteries, a short helicoidal antenna, a manometer and low density stuffing for flotation. The antenna is efficient against waves and the manometer stops emission during dives, restoring it a few meters below the surface (i.e. a few seconds before the Bp surfaces). Previous observations of Bp in the Mediterranean had shown that they often take very short surface breaks between dives, too short to complete an Argos emission cycle.

The PTT was not in contact with the whale’s body: a 12 m line tied to the dorsal fin of the 20 m long animal, dragged the Argos device. The attachment system between the Bp and the PTT had been designed to respect the animal’s physiology. If fixed to its flank, the fragile skin would not have supported a 2.5 kg buoy, which would have caused a wound. To avoid that, the attachment setup [6] [12], used a Kevlar line between the dorsal fin and the Argos buoy.

Finding a Bp in natural conditions, approaching the animal and attaching the Kevlar line to its dorsal fin, required three years of training and trials. It was achieved by Carli Viale on 22 September 1991. The Argos buoy transmitted its position to the satellites during 42 days.

## 3. Data Mining and Analysis

Modern technologies allow a new look at these old data. The digital data from 1991 were irrecoverable, but the paper dumps remained readable. They were scanned and redigitized using optical character recognition (OCR).

Several track points were erroneous and have been removed. We used two criteria to identify errors: the Signal/Noise ratio given by CLS-Argos for each point, and unfavourable positions of the Argos satellites. The precision given by the Argos Doppler tracking is poor when the satellite-to-whale distance passes through a minimum (satellite at zenith for example) and, for different reasons, when the satellite is low above the horizon. These imprecise points added bogus spikes and high speed segments. A few of them remain, e.g. on 3 and 18 Oct. (thereafter, the year: 1991, is implicit).

Argos yields geographical coordinates (longitude, latitude), which we translated into respective distances (in km) from the equator and from the Greenwich meridian, yielding one point (two coordinates) on the map. Then, using pairs of points, we derived segments (four coordinates) and their average speed, and from each couple of segments (i.e. three consecutive points or six coordinates) we derived a signed angle in the trajectory. We integrated these angles to detect loops in the trajectory, see fig.5.

To each segment we associated the local times of sunrise and sunset, in order to study the effect of the daily cycle on the animal’s behaviour. We also investigated the causes of non reception of signals: satellite not high enough above the local horizon, or whale submerged.

## 4. Results

The whale trajectory from 22 Sept. to 02 Nov. has a cumulated segments length of 1317 km. The actual trajectory is longer, being fractal and undersampled [11].

The Argos satellites received between 2 and 15 ‘messages’ per day. Their high number per day was a surprise, as the Argos buoy cannot send messages when the Bp is submerged: it appears that this Bp was more often near surface when left alone than when in the vicinity of human observers.

The transit duration of an Argos satellite above the whale’s horizon lasted from 20 to 800 s. Our prior observations in the Mediterranean showed Bp dives lasting from 240 to 900 s, most frequently 450 to 500 s. Our expectations were pessimistic. Fig.1 displays the number of messages received and the duration of satellite transits over the Bp as a function of time (modulo 24h). Because there were only two satellites, the Argos coverage is variable and incomplete. The fin whale was not covered by the satellites during the following intervals each day: 20h50–01h30; 05h00–06h00; 10h00–12h00; 15h50–16h30.

**Figure 1:**
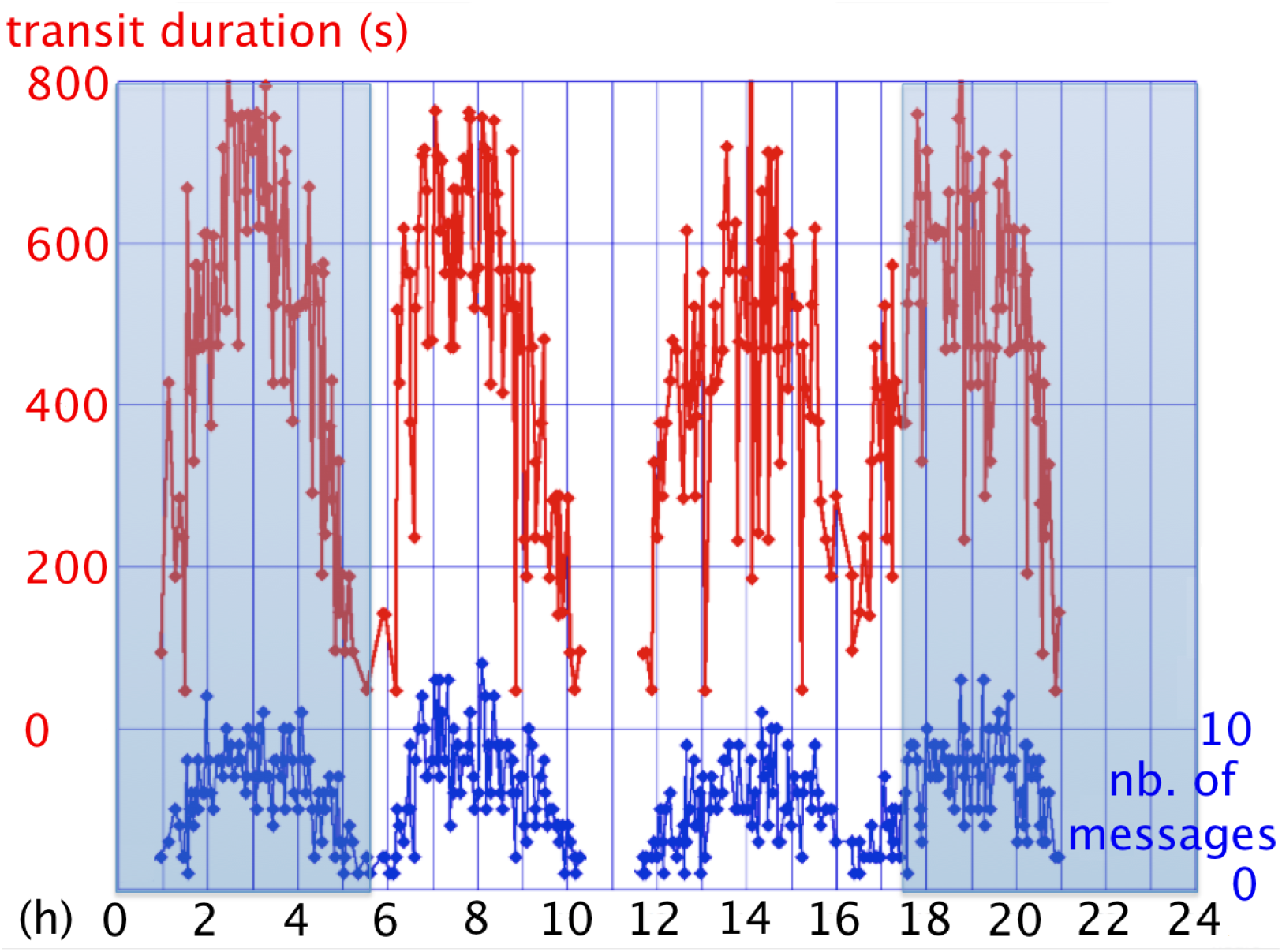
Satellite coverage of the whale as a function of local time (hours 0 to 24). In red: transit durations of the Argos satellites above the local horizon; in blue: number of satellites messages in a given one-hour interval, cumulated over the 42 days tracking.

From our revised study of: the satellite schedules; the message lengths and transit durations; the positions of the Bp; the local times of sunrise and sunset, we have deduced some aspects of the whale’s behaviour. These are synthesized in fig.2, which shows the track superimposed on a GoogleEarth map of the area.

**Figure 2:**
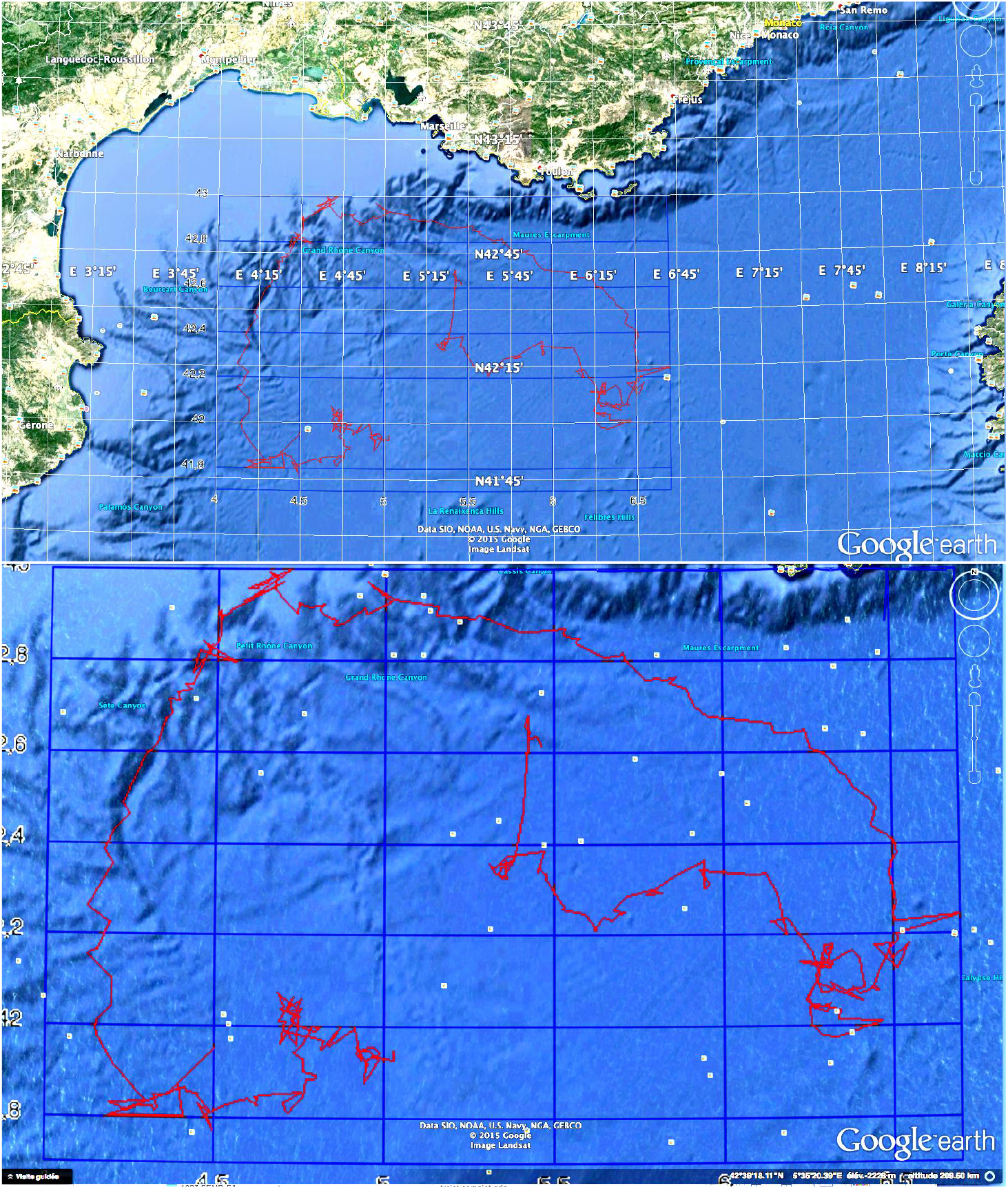
Track of the fin whale superimposed onto the corresponding Google Earth map of the north-western Mediterranean Sea (Corsican coast at right; French coast, Toulon, Marseilles, at top; Spanish coast, Girona at left). The bottom figure is an enlargement, better showing correlations between the whale’s track and submarine canyons or cliffs. Note the match of the track to the underlying submarine topography, the marine circulation, and sources of food, such as the upwelling zone south of the Gulf of Lion.

Unlike that of a drifting buoy, the whale track shows active adaptation (dubbed ‘contagious’) to the marine environment and food resources. The plankton and krill are not distributed randomly, being concentrated in areas where the joint action of marine circulation, winds, temperature and submarine topography enrich the sea in nutritive salts [7] [12] [14] [15] [16]. The shape of this track suggests several biomass-rich zones in the Mediterranean.

These data on the Mediterranean ecosystem had been collected previously by participating in the international oceanographic mission “Western Mediterranean Circulation Experiment” [13] [15]; here they explain the Bp speed modulations observed: fast through food-sparse zones (e.g. in the Catalan stream from 18 to 21 Oct.) or slow in non-circulating water masses (22 Oct. to early Nov.), where long chains for biomass production have time to develop from phytoplankton: → phytophagous zooplankton → phyto- and zoophagous zooplankton → macroplankton, such as Euphausiacea, forming swarms hunted by the Bp.

## 5. Discussion

These inertial fluctuations correspond to an oceanological phenomenon due to Coriolis force, observed with the drifting buoys used to study superficial circulations [9] [17]; they reveal rotations of sea-water masses, within which the buoy (or whale in our case) is ‘inert’. These loops would not cause a displacement of a whale relative to a surveillance boat: both drifting in the same water mass.

Our reprocessing of the Argos data revealed 21 loops in the whale’s track. To detect these loops we integrated the angles between consecutive segments of the trajectory, as explained in sect.3. The resulting curve is plotted in fig.5. It turned out to be an increasing function of time, except in the central part of the graph where the whale was in the main Mediterranean circulation. The strength of sea circulation prevents the Coriolis loops from closing, which results into a cycloidal trajectory. The undersampling of the Argos coverage turns it into a ‘sawtooth’ shape.

From 15 to 18 Oct. there is a short ‘loopy’ phase, when the whale is above the Bourcart canyon. It is localised at a quasi-constant position: 10 times on 16 Oct. and 10 times again on 17 Oct. in the same 8×8 km area; a total of 105 Argos messages in 35 hours, which provide information on the whale’s behaviour: 15 messages during daytime on 17 Oct. and 90 messages, by clusters of 6 or 7 during the adjacent nights. The last message clusters received before these mornings are at 17 Oct. 03:54 and 18 Oct. 02:03; by the time of the following satellite coverage the whale had resumed its activity. As a consequence, there were three times more messages when the whale was surfacing in inertia at night than during its periods of activity.

The third part of graph in fig.5 rises again, accounting for six turns, and corresponding to the ‘complex’ end of the track, from 26 Oct. to the last recorded position 02 Nov. Overall, we detected 26 phases of inertia, which are periodic with a 24h period. We interpret the ‘free floating’ periods during which the whale shows no autonomous mobility, being due to the animal being at muscular rest, a state that seems to represent sleep.

To ascertain the link between repeated inertial loops and sleep phases, it has to be shown that they occur every night. These loops are conspicuous only in the parts of the trajectory that are drawn in slow water masses (see fig.4), such as near 01 Oct., away from the northwestern Mediterranean circulation. They do not show up in areas with circulating water masses, where, like drifting buoys, the Bp trajectory presents broken line segments [9][17], for example from Oct.11 to 15, in the ‘Provençal current’ and from Oct. 18 to 22 in the ‘Catalan current’. The animal is taking advantage of, rather than being subjected to, this circulation: on 22 Oct. it leaves it and moves towards rich zones, possibly guided by vocalizations of other animals. We have evidenced elsewhere that the speed of the Bp at night between 20h30 and 01h50 appears to be precisely in correspondance with the current, whereas during daytime the Bp is tracked at speeds twice faster than the current. This allows to distinguish ‘inertial’ phases at night from ‘active’ phases during the day [23].

In the upper right part of the track leading North-West, 06 to 14 Oct., (fig.2) the Bp is in a region where intermittent currents due to cold North winds (e.g. mistral) cause the colder, denser surface water to sink. This in turn causes an upwelling of ‘intermediate water’, rich in nutriments, into the euphotic layer. Such local turbulences are sources of nutriment enrichment, leading to a possible increase in biomass production, including krill. This food supply in turn would be taken advantage of by the wale, as suggested by the wiggles (variable segment lengths, speed and direction) in this part of its track (fig.4). The speed varies because the whale travels faster through food-poor regions. One can see (fig.3) that night-time segments are quasi similar, varying with the stream speed. Short day-time segments indicate a diving activity that ceased one hour before sunset, and/or a direction of swim northwards, opposite to the current.

**Figure 3:**
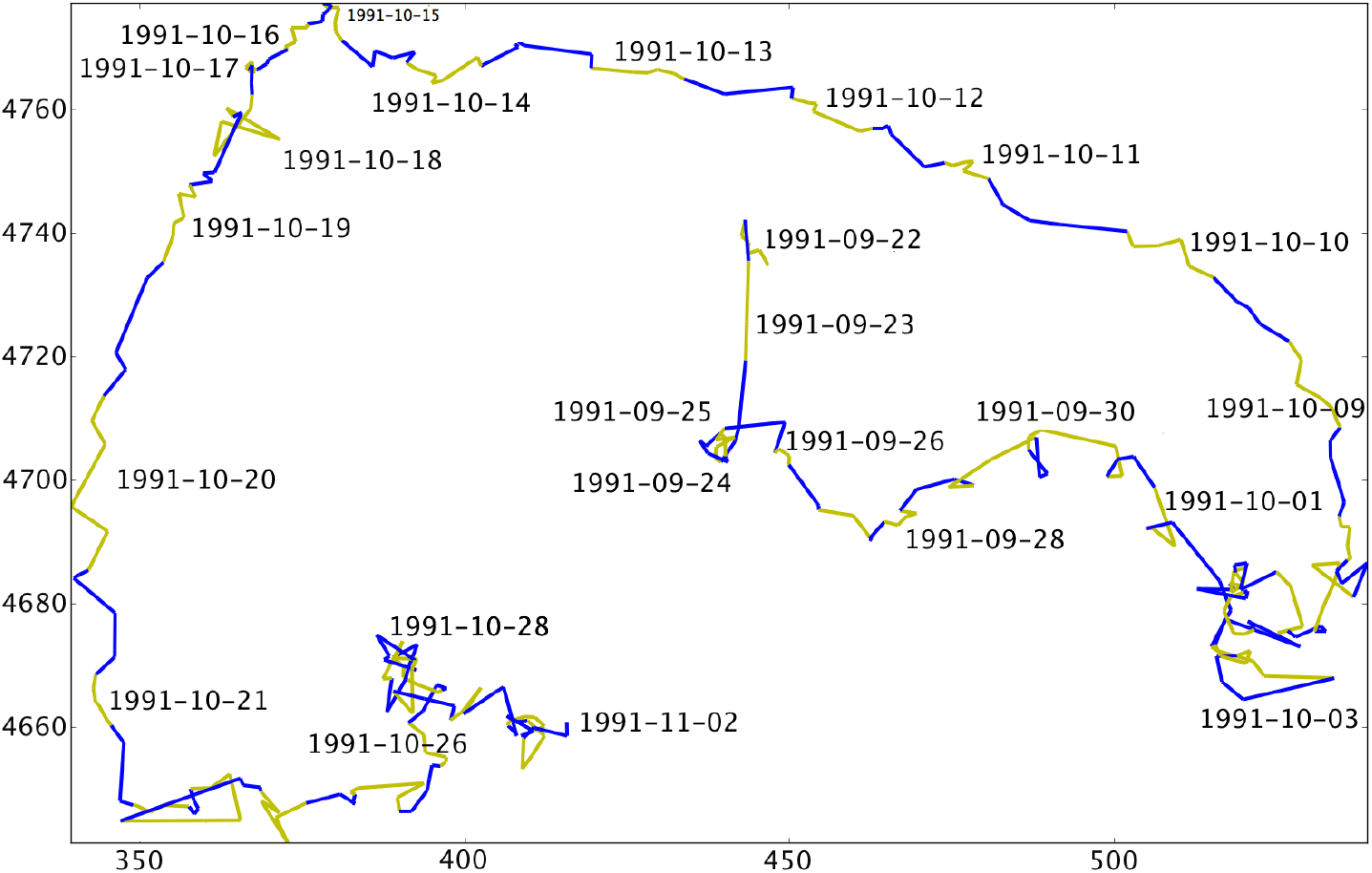
Day-night alternation of the Bp between consecutive localisation points, colour-coded: in blue are the track segments starting during nighttime, and in olive during daytime. Scales in km: vertically from the equator; horizontally from the Greenwich meridian.

The part of the track from 22 to 25 Oct. remains in an area interpreted as being rich in trophic resources because the Bp track becomes complex and slow. Around 28 Oct. the track points are numerous and frequent: the Bp emitter is more often detectable as the animal probably dives to feed and rests longer on the sea surface. The six inertial loops (plus one night in circulating water) in the last part of the track are hard to see, but their presence is confirmed by the integrated angles procedure (fig.5).

The track features 26 inertial loops plus 15 inertial phases in circulating waters during the 42 days and 41 nights it covers (the first day and last night were not covered entirely). It appears that our Bp had a sleep phase every night.

On fig.3, the majority of the loops and related segments occur between two hours after sunset and the middle of the night. These daily periods after sunset, when the Bp is inactive, correspond to the rise and dispersion of plankton over the sea surface. Our previous work had shown that Bps dive for feeding only into concentrated biomass volumes: swarms formed during vertical nycthemeral migrations of the krill. These krill swarms remain developed at 50-75 m depths, between one and one and a half an hour before sunset. Then they disperse over the euphotic layer, where they consume the plankton matured during the day [15][16] and present no interest to the Bp, which stops diving one hour before sunset. The descent of the krill begins one hour before sunrise; in the following hour biomass accumulates at the usual depths and the whales start diving again.

The inertial loops have allowed us to estimate the duration and time of sleep phases: around 6 to 8 hours, starting after sunset, whereas feeding activity stops one hour before. These are estimated with limited precision, as the two satellites of the Argos system during these early years of global tracking did not cover the this part of the Mediterranean Sea between 20:50 and 01:15.

## 6. Conclusion

The tracking of the whale in 1991 had originally allowed the detection of five cycles of activity followed by rest. The hypothesis of a sleep phase was suggested, but only for a few days in the tracking record. Rest periods were revealed by the α-shaped loops of the whale track. In addition, the Argos tracking indicated that the animal was at the sea surface (not diving) during those periods. A cetacean’s sleep implies two conditions: ability to breathe, hence at or close to the surface, and no displacement relative to the surrounding water [18].

In the present work we retrieved and reprocessed the 29 year-old data. We superimposed the track onto a Google Earth map (fig.2), on which the path of our tagged fin whale appears coherent with its marine environment. For each segment of the track we calculated an average speed (fig.4) and correlated this with the local times of sunrise and sunset (fig.3), to study the effect of day/night cycles on the whale’s behaviour. The length and speed of the track segments recorded during day-time vary as a consequence of diving for food and its availability, whereas night-time segments are more constant in length, varying in accordance with the speed of the stream in which the Bp is swimming (or floating when in a phase of inertia). The graph in Fig.5 is very different from one that would be obtained from a random trajectory; it reveals Coriolis loops in the Bp track every night, except when currents prevent them from closing.

**Figure 4:**
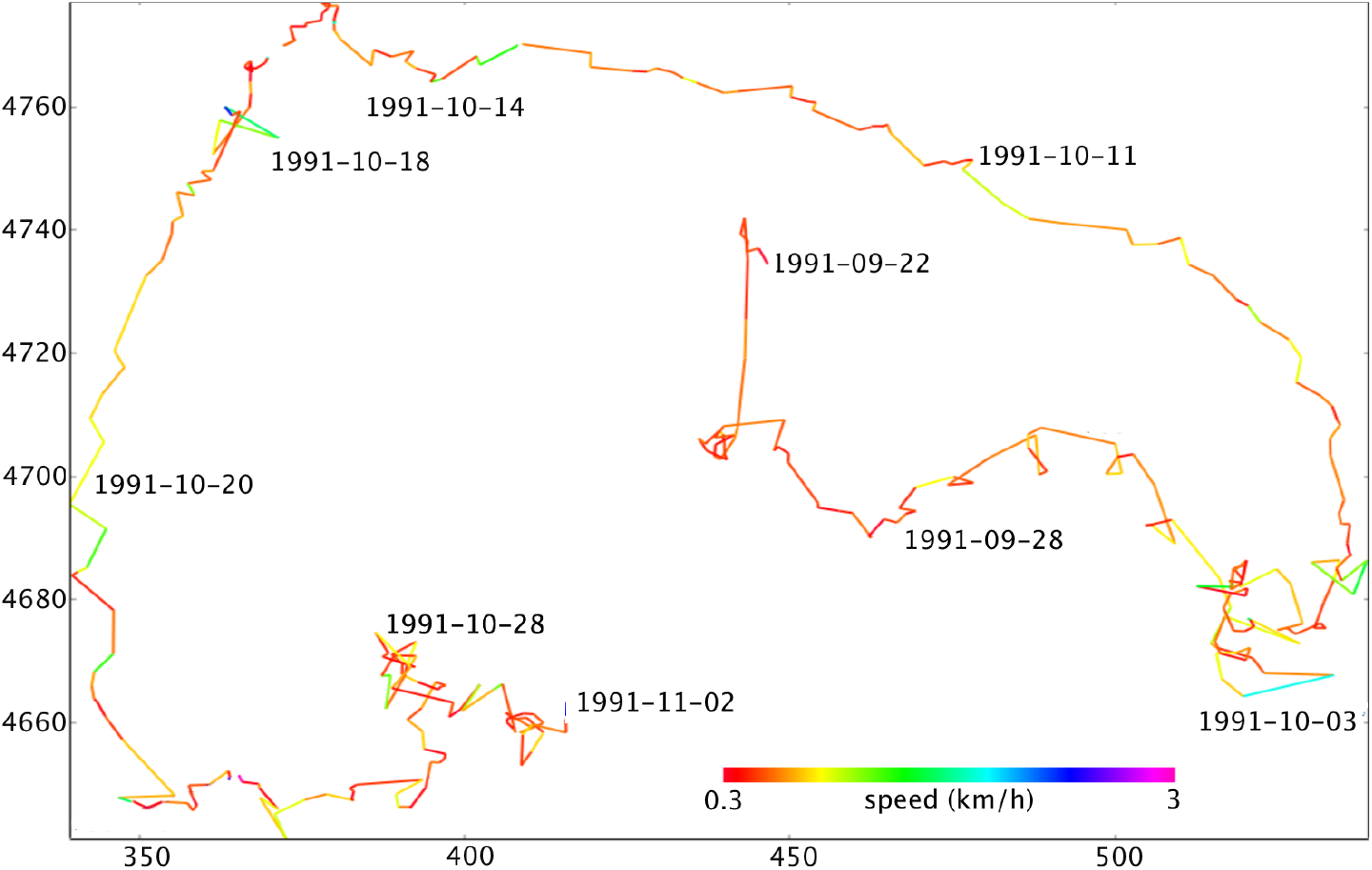
Average speed of the Bp between consecutive localisation points, colour-coded from red: ≃ 0.3km/h to blue: ≃ 3km/h. Scales in km: vertically from the equator; horizontally from the Greenwich meridian. The speed is high, for example, when the animal is in the Catalan stream (descending on the left side of the plot). Two high speed segments (in blue, tagged 1991-10-03 and 1991-10-18) are due to erroneous points in the Argos tracking: poor precision when the satellite is close to zenith.

**Figure 5:**
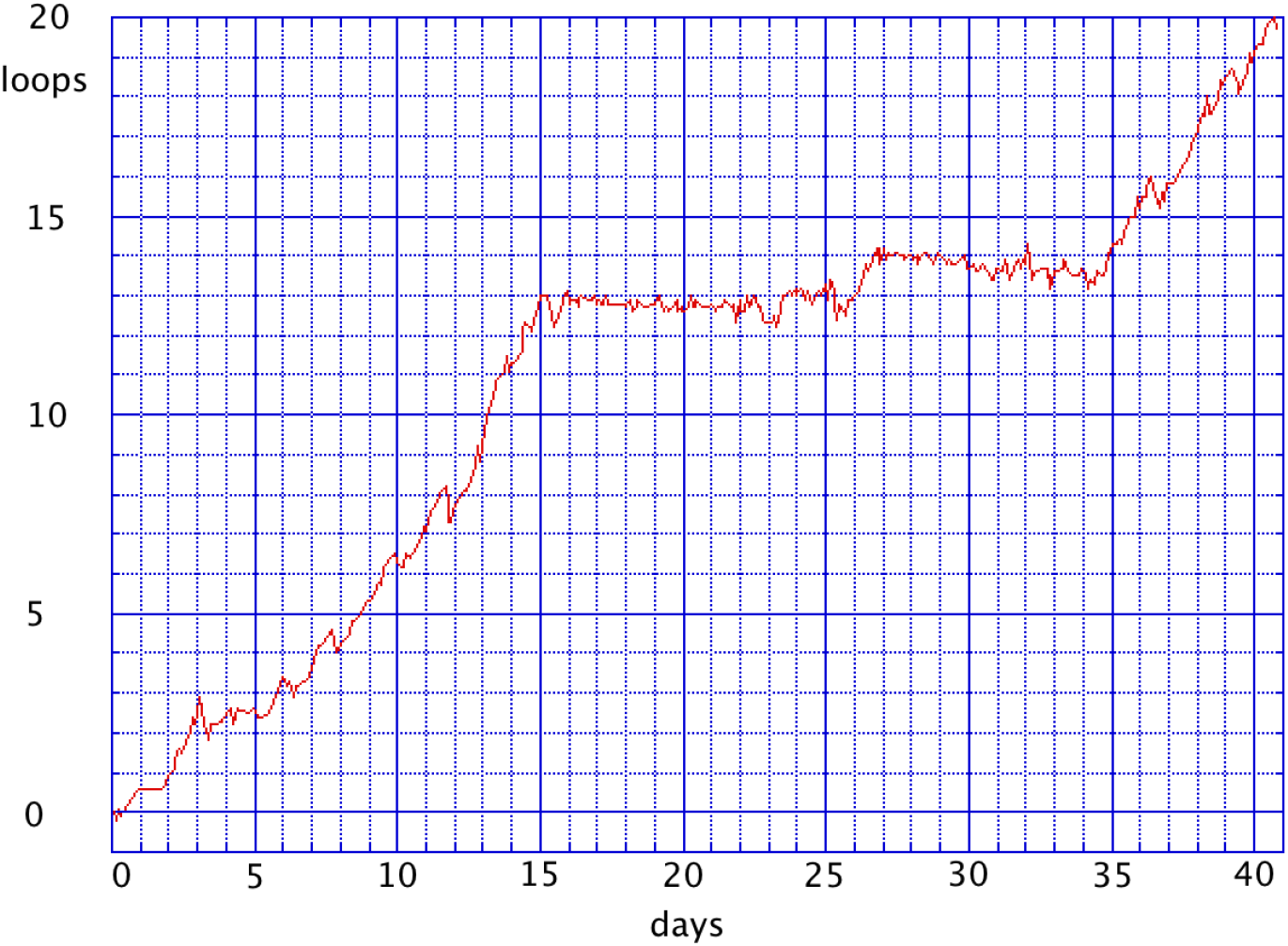
Graph showing the number of loops detected in the whale’s track as a function of time. Ordinate: cumulated angles between consecutive track segments (number of loops since origin, 360° = one loop). This procedure allows to detects loops that do not appear clearly to the eye in the plot of the Bp track. Abscissa: time (in days) since first day of tracking. The fact that this curve is increasing with time means that all loops are executed in the same direction: clockwise, except in the central part of the graph from 6 to 26 Oct. (days 15 to 35).

The 20 m long whale that we tracked behaved like a ship without a harbour: it did not need to stop over for food or fresh water. We have shown [19] [20], that a Bp desalinates its internal physiology by means of a pleated surface of the germinal epidermal layer and the dermis in contact with a vast layer of blood capillaries of the hypodermis. It ‘sweats’ salt crystals by regularly peeling off its superficial epidermal layer.

This ‘ship’ had another problem to solve: sleep. Sleep in cetaceans has been shown to be uni-hemispheric [21], here we simply deduce a sleep cycle for Bp from our data on its behaviour, we make no further assertions regarding its cerebral sleep mechanisms. Anecdotal accounts exist, observed (or even harpooned) by ships [22]. But were these animals asleep, or sick, or dead? This harbourless whale has indeed places to rest in the marine environment: moored to its water each night. The measured speeds of the Bp during these nighttime segments, when in circulating waters and when they are identical to the water current, indicate that inertial phases are present. Such phases of total inertia, with the absence of autonomous movement, usually start one or two hours after sunset. They correspond to a ‘rest’ of the cerebral commands for motion and at least partly accomplish the functions of sleep. This indicates the regular sleep cycle of a whale lasting approximately 10 hours each night.

## 7. Acknowledgements

We pay tribute to R. Chesselet, founder and head of INSU at CNRS, who enthusiasticly supported us when we started the project.

This project was financed by: CNRS Aide n° 34915 AD12, Laboratoire d’écologie des cétacés; Université Paris VI (ERA 715, Prof. P. Nival, Villefranche sur mer); Université de Corse (Faculté des sciences), Région Corse, Armée de Terre, Légion étrangère (2ème REP de Corte); Université Paul Sabatier (Toulouse); and private funds.

Our thanks to: “Comité Inter-Régional Méditerranéen” for the allocation of vessels and means at sea of CNRS; Commission Nationale CNRS-IFREMER for the allocation of vessels for remote missions (Noroit, Suroit, Charcot); J.J. Pesando for his engineering work; P. Braconnier and J. Roqueferre for their courageous help to C. Viale in the whale approach; N. Terris, I. Palazzoli, J.P. Frodello and D. Mouillot, who participated to the project; the crews of N/O Korotnef and Antedon, and particularly the seamen of N/O ‘Catherine Laurence’ who proposed to stay at sea on the evening of 21 Sept., thus leading to the success in tagging the whale the next morning; the weather services of Meteo France (CMS Lannion) for their maps of thermal fronts; MM. Taillade, Gros and Ortega of CLS Argos (Centre de localisation par satellite Argos) in Toulouse, France; L. Masson, who scanned and ‘OCRd’ the old Argos printouts; all the sailors, CNRS engineers, and volunteers who contributed to the success of this Argocet mission; and Anne & Julien Viale, who have supported it for so many years.

